# Blocker-SELEX: A Structure-guided Strategy for Developing Inhibitory Aptamers Disrupting Undruggable Transcription Factor Interactions

**DOI:** 10.1101/2024.01.09.574928

**Authors:** Tongqing Li, Xueying Liu, Sheyu Zhang, Yu Hou, Yuchao Zhang, Guoyan Luo, Xun Zhu, Yanxin Tao, Mengyang Fan, Chulin Sha, Ailan Lin, Jingjing Qin, Weichang Chen, Ting Fu, Yong Wei, Qin Wu, Weihong Tan

**Affiliations:** Hangzhou Institute of Medicine, Chinese Academy of Sciences, Hangzhou, China; School of Pharmacy, Zhejiang University of Technology, Hangzhou, China; Shanghai Institute of Material Medica, Chinese Academy of Sciences, Shanghai, China; School of Life Sciences, Tianjin University, Tianjin, China; Hangzhou Institute for Advanced Study, University of Chinese Academy of Sciences, Hangzhou, China

**Author notes:** Corresponding authors: Weihong Tan,; Yong Wei,; Qin Wu. These authors contributed equally (Tongqing Li, Xueying Liu).

**Keywords:** Inhibitory Aptamer, Blocker-SELEX, Protein-protein interactions, Undruggable

## Abstract

Despite the well-established significance of transcription factors (TFs) in pathogenesis, their utilization as pharmacological targets has been limited by the inherent challenges associated with modulating their protein-protein and protein-DNA interactions. The lack of defined small-molecule binding pockets and the nuclear localization of TFs do not favor the use of small-molecule inhibitors, or neutral antibodies, in blocking TF interactions. Aptamers are short oligonucleotides exhibiting high affinity and specificity for a diverse range of targets. Large molecular weights, expansive blocking surfaces and efficient cellular internalization make aptamers a compelling molecular tool for use as traditional TF interaction modulators. Here, we report a structure-guided design strategy called Blocker-SELEX to develop inhibitory aptamers (iAptamer) that selectively block TF interactions. Our approach led to the discovery of iAptamers that cooperatively disrupts SCAF4/SCAF8-RNA Polymerase II (RNAP2) interactions, thereby dysregulating RNAP2-dependent gene expression and splicing and, in turn, leading to the impairment of cell proliferation. This approach was further applied to develop iAptamers to efficiently block WDR5-MYC interaction with a nexus in cancer. Taken together, our study highlights the potential of Blocker-SELEX in developing iAptamers that effectively disrupt potentially pathogenic TF interactions with attendant implications for iAptamers as chemical tools for use in the study of biological functions of TF interactions, but also for potential use in nucleic acids drug discovery.

## Introduction

Transcription factors (TFs) represent a pivotal protein class that orchestrates gene transcription, exerting profound influence over cellular identity and fate^1–3^. TF activity is finely tuned through intricate, dynamic interactions with DNA regulatory elements and TF-binding proteins^4–7^. Dysregulation of TF activity has been implicated in a plethora of human diseases, encompassing cancer, infectious diseases, and neurodegenerative disorders^8^. Consequently, targeted modulation of TF interactions has emerged as a highly promising therapeutic strategy^9–13^.

A wide range of molecules, including small molecules, peptide mimics, and antibodies, have been employed as modulators of protein-protein interactions^12^. Small molecules, a traditional approach, are utilized to block interactions that involve proteins with well-defined ligand-binding sites^14^; accordingly, small molecules have been successfully used in blocking well-folded TF interactions. A successful example is the development of small molecules capable of disrupting p53–MDM2 (mouse double minute 2) protein interactions, a target in the development of antitumor drugs, such as Idasanutlin and AMG232, which have been designed and subjected to clinical trials (NCT02545283, NCT02110355, et al.)^15,16^. However, most TF interactions are intrinsically disordered or lacking in well-folded small-molecule binding pockets, posing challenges for small molecule-based interventions. Peptide mimics, derived from peptide fragments involved in protein interfaces, also serve as modulators of protein interactions^17,18^. For example, OmoMYC and STRs can disrupt MYC-Ebox interactions, previously considered to be an impossible mission^19–21^. However, the clinical translation of peptide mimics has been hampered by certain intrinsic limitations, including intracellular localization, targeting tissue specificity, and pharmacological potency^22^. Recent advancements have witnessed remarkable progress in developing antibodies as modulators of protein-protein interactions^23–25^. Antibodies blocking PD-1 and PD-L1 interaction, such as pembrolizumab and Opdivo^®^ (nivolumab), have exhibited remarkable efficacy in cancer treatment^26,27^. Nonetheless, antibodies face limitations in targeting intracellular entities owing to their inherent inability to traverse cellular membranes^12,28^. Consequently, interest is growing in identifying and developing alternative classes of modulators capable of targeting intracellular TF interactions.

Aptamers are short oligonucleotides demonstrating remarkable specificity and high affinity towards a diverse repertoire of targets, encompassing proteins, sugars, phospholipids, and cell metabolites. Their versatility has attracted significant attention based on their potential applications in therapy, diagnostics, bioimaging, and drug delivery^29,30^. Notably, the U.S. Food and Drug Administration (FDA) has granted authorization for three aptamer drugs, namely Pegaptanib, Apc001PE and Zimura (avacincaptad pegol), all serving as potent modulators of protein-protein interactions. Pegaptanib, approved in 2005 for the treatment of age-related macular degeneration, exerts its therapeutic effect by selectively binding to the surface of vascular endothelial growth factor (VEGF), thereby abrogating its interaction with the cellular receptor VEGFR^31^. Apc001PE, authorized as an orphan drug in 2019 for osteogenesis imperfecta, functions by disrupting the interaction between sclerostin (SOST) and low-density lipoprotein receptor-related protein 5/6 (LRP5/6)^32–34^. Avacincaptad pegol, approved in 2023 for geographic atrophy, inhibits the interaction between Complement C5 and its receptor C5R^35,36^. These achievements underscore the therapeutic potential of aptamers that specifically target protein interactions^30^. However, it is crucial to note that most reported aptamers primarily focus on interactions involving cell membrane proteins. Several natural prokaryotic and eukaryotic examples of RNA aptamers that modulate the activity of TFs can be cited. However, only a few unnatural aptamers have been selected to inhibit TF-DNA interactions, including NF-κB, TBP, HSF1, and RUNX1^37^, owing to the lack of selection method for aptamers targeting TF interactions.

To solve this dilemma, we herein present a structure-guided rational design strategy, termed Blocker-SELEX, for the development of inhibitory aptamers (iAptamers) that effectively disrupt TF protein-protein interactions. Through a structure-guided screening of a single-stranded DNA (ssDNA) library against the interfaces derived from TF complex structures, we identified and validated a lead sequence with demonstrated target binding and competitive properties. We further optimized the lead sequence by incorporating supporting nucleotides to optimize affinity and competition efficacy. Using this strategy, we successfully developed iAptamers that inhibit the interaction between SCAF4/SCAF8 and RNAP2 proteins. Treatment of tumor cells with selected iAptamers altered the profile of cell transcripts, dysregulated alternative RNA splicing, ultimately reduced cell proliferation, and increased apoptosis. The versatility of our approach was further validated by successfully developing iAptamers targeting the intrinsically disordered oncogenic MYC protein that interacts with WDR5^20,38^. Altogether, our study has demonstrated a structure-guided design strategy for the development of iAptamer tools that specifically block intracellular TF interactions, highlighting their promising implications in investigating the biological functions of TF interactions. Given the rapid advancements in nucleic acid delivery techniques, these iAptamers hold promise for nucleic acid drug therapeutics.

## Results

### Development of Blocker-SELEX pipeline

To design aptamers able to block protein-protein interactions, we proposed a strategy consisting of four main procedures: ⍰ structure-guided virtual screening, ⍰ lead sequence validation, ⍰ iterations of sequence optimization, and ⍰ iAptamer validation (Fig. 1a). Structure-based virtual screening has been widely used for lead compound discovery in the early stage of small-molecule drug development. This strategy utilizes the three-dimensional structure of target proteins to dock a collection of chemical molecules into the desired binding site, and a subset of compounds with favorable binding scores is selected for further functional evaluation^39^. While recent attempts have integrated virtual screening with aptamer SELEX, validation of the results can still be challenged by limitations in accurately predicting 3D structures of aptamers in the library pool^40–42^.

**Figure 1:**
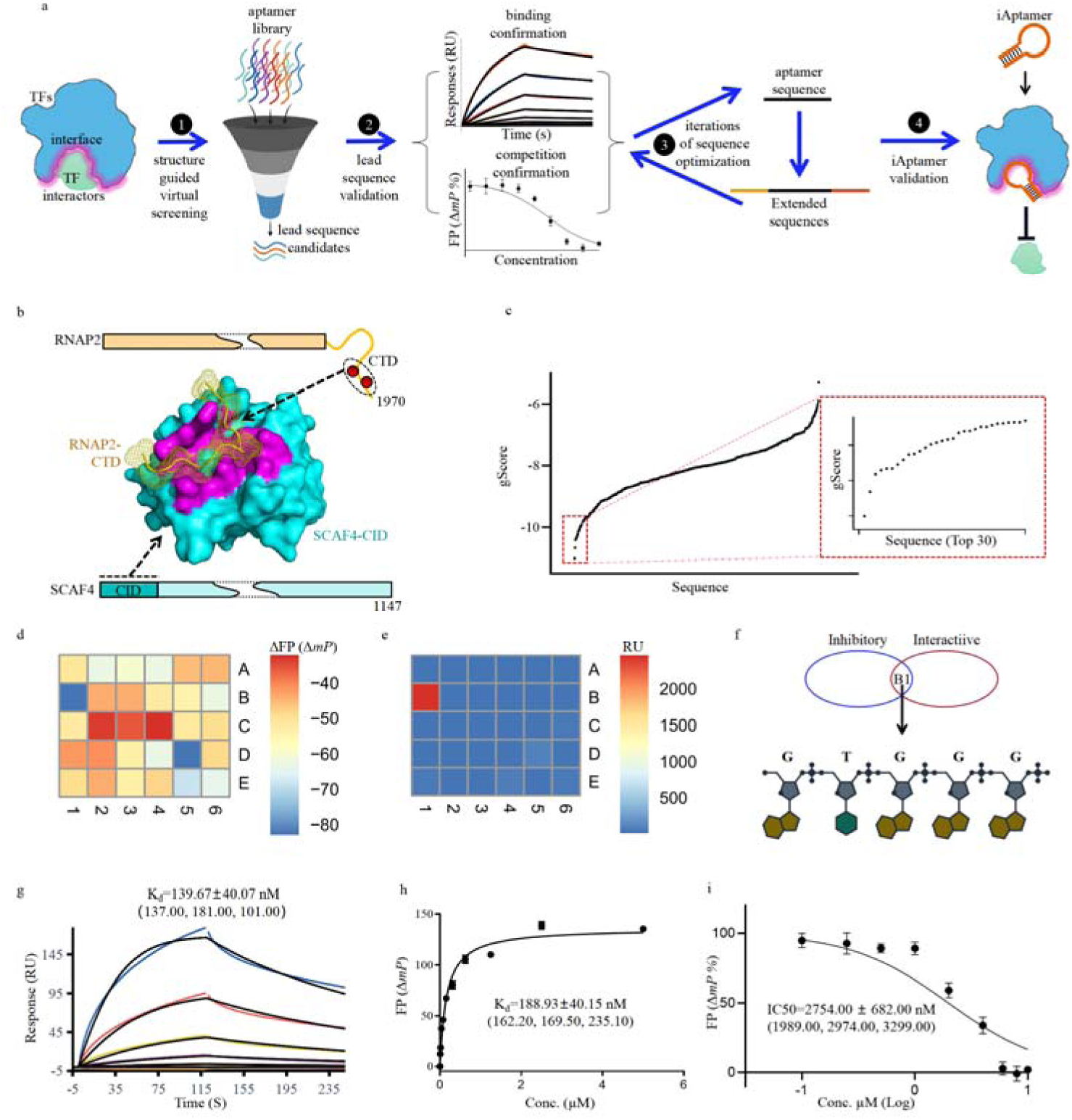
Identification of the lead sequence binding to SCAF4 and blocking its interaction with RNAP2. **a**) Schematic representation of structure-based design strategy for aptamer inhibitors. **b**) Surface representation of SCAF4-RNAP2 complex structure re-illustrated from PDB 6XKB. SCAF4 protein is depicted in cyan and RNAP2 in yellow, while the interface is colored in pink. **c**) Docking scores of 5-nt sequences to SCAF4 from the Glide pipeline. The top 30 sequences were enlarged in the inserted panel. **d**) Inhibitory effects of top 30 sequences to SCAF4 and RNAP2 interaction were determined by competitive fluorescent polarization assay. **e**) Direct interactions of top 30 sequences to SCAF4 were determined by SPR spectroscopy. Proteins were exposed to 10 μM of 30 top scored 5-nt sequences. RU, response units. **f**) Selection of DNA sequences with both binding affinity and inhibitory effects. Chemical structure of the identified lead sequence was inserted. **g**) Binding of SCAF4_LS to immobilized SCAF4 by standard kinetics SPR. This assay was performed with three replicates. Average affinity value was 139.67 ± 40.07 nM. Affinities of each replicate was indicated in the bracket. **h**) Fluorescence polarization measurements where 50 nM FAM-labeled SCAF4_LS was titrated with increasing amounts of SCAF4 proteins. The affinity was determined as 188.93 ± 40.15 nM using the FP (ΔmP) signals with three biological replicates. Affinities of each replicate are indicated in the bracket. The non-linear fit model and the one-site specific binding to fluorescent population ratio were used for these calculations at the respective aptamer concentrations. **i**) Dose-response curves of SCAF4_LS showing the inhibition of RNAP2 interaction with SCAF4. The half-maximal inhibitory concentrations were determined from three independent experiments, each containing three independent replicates. IC50 values were calculated using GraphPad Prism software (version 7) with the Log(inhibitor) vs. normalized response model. IC50 values of all experiments are indicated in the bracket (n=3, mean ± SD).

To address this issue, we propose a screening methodology grounded in a library of short DNA sequences characterized by relatively reduced structural complexity. Extensive structural studies have revealed that aptamer recognition of target molecules often relies on a limited number of key bases, such as the aptamers for Thrombin, GRK2, and Hfq^43–45^. Thus, a library of 1024 (4^5^) single-stranded DNA (ssDNA) sequences with 5 nucleotides was generated for initial virtual screening. The chemical structures of these sequences were created using JChem software (http://www.chemaxon.com) and optimized using the LigPrep tool^46^ to achieve optimal chemical structures. The binding potential of each DNA sequence to the target protein was calculated using GlideScore, an empirical scoring function that approximates ligand binding free energy^47,48^. Binding affinity and blocking efficacy of the top-scoring sequences with highest binding potentials were determined by both surface plasmon resonance (SPR) and competitive fluorescence polarization assay. Those DNA sequences exhibiting a dual capacity, i.e., both target binding and concurrent inhibition of interaction, were identified as lead sequences serving for subsequent optimization.

Traditional aptamers basically consist of two distinct regions: core regions and scaffold regions^49–51^. Scaffold regions serve as a framework for the core regions, thereby heightening their specificity and affinity towards targeted molecules. In light of this insight, we set out to strengthen the affinity and inhibitory efficacy of the identified lead sequence by iteratively adding nucleic acid bases to the lead sequence. In each iteration, a set of three bases was appended to the terminal end of the lead sequence, resulting in the generation of 64 (4^3^) optimized sequences. The rationale underlying the incorporation of three bases in each iteration is rooted in practical considerations. First, the resulting 64 optimized sequences can be handled on a single 96-well plate to determine their affinities to the targeted TFs and their increased competitive capabilities in blocking TF interactions. Second, this strategic approach aligns with cost-effective concerns in nucleic acid synthesis. Upon achieving the desired levels of binding affinity and competitive potency, the fine-tuned aptamers are officially designated as iAptamers, marking the culmination of this optimization process.

### Identification of a Lead Sequence Targeting SCAF4-RNAP2 Interaction using Blocker-SELEX

To investigate the efficacy of the proposed Blocker-SELEX pipeline in generating iAptamers capable of blocking TF protein-protein interactions, we selected the interaction between SCAF4 and RNAP2 as our target. SCAF4, a TF that interacts with the C-terminal domain (CTD) of RNAP2, plays a crucial role in regulating its transcriptional activity. Dysregulation of SCAF4-RNAP2 interaction has been implicated in erroneous poly-A modification, aberrant transcriptional termination, and cell proliferation arrest^52^. A previous study of SCAF4-RNAP II interaction has revealed that it is an undruggable pocket, which underscores the challenge in devising traditional pharmacological strategies (Fig. 1b; Supplementary Fig. 1a)^53^. Consequently, the absence of specifically designed chemical tools has hindered in-depth exploration of the precise biological function associated with this interaction. Therefore, we aimed to design iAptamers that could effectively target the interface between SCAF4 and RNAP2, thereby disrupting their interaction and impeding their functional partnership.

To prepare the docking receptor, the structure of SCAF4 was extracted from the SCAF4-RNAP2 complex (PDB code: 6XKB, chain A). Specifically, we selected the protein-protein interaction (PPI) interface of SCAF4 that recognizes the RNAPII-CTD domain as the docking target (Fig. 1b). 4^5^ ssDNA molecules from the library were docked against the prepared receptor of SCAF4 to evaluate the binding potentials using GlideScore (Fig. 1c). Among them, the top 30 ssDNA sequences with the highest Glide scores were selected for synthesis and subsequent experimental validation to assess their inhibitory efficacies against SCAF4-RNAP2 interaction, as well as their binding to SCAF4, using SPR. To investigate the potential inhibitory effects of these sequences on SCAF4-RNAP2 interaction, we established a competitive screening assay based on fluorescence polarization. Initially, the interaction between SCAF4 and RNAP2-CTD was captured with a phosphorylated CTD peptide (pS2pS5) of RNAP2 labeled with an NH2 terminal FITC group for polarization measurements. The affinity between phosphorylated CTD and SCAF4 was determined as 365.60 nM (Supplementary Fig. 1b). For the competition screening, FP signals (> 60 ΔmP) were generated by combining 2 µM SCAF4-CID with 40 nM FITC-labeled CTD peptide solutions. Reductions in FP signals were observed upon the addition of the top-scoring sequences. Notably, sequences 5’-GTGGG-3’ and 5’-CTGGG-3’ exhibited significant signal reduction, suggesting their competitive roles in blocking SCAF4-RNAP2 interactions (Fig. 1d). Concurrently, we investigated the interactions of these top-scoring sequences with SCAF4 protein. To accomplish this, purified SCAF4 protein was immobilized onto a CM5 chip, and 30 aptamer candidates were assessed for their binding specificity by a single injection at a concentration of 10 µM for screening (Fig. 1e). As summarized in Figure 1e, ssDNA 5’-GTGGG-3’ demonstrated significant binding to SCAF4 with a response unit exceeding 2000 (RU), while the remaining sequences displayed only weak, or negligible, responses. Based on these findings, ssDNA 5’-GTGGG-3’ with both binding affinity and inhibitory effects was selected for further determination (Fig. 1f). The binding affinity of ssDNA 5’-GTGGG-3’ to SCAF4 protein was further determined to be 139.67 ± 40.07 nM by SPR (Fig. 1g; Supplementary Fig. 1c). Consistently, a fluorescence polarization assay verified its affinity to be 188.93 ± 40.15 nM (Fig. 1h; Supplementary Fig. 1d). Finally, the inhibitory capacity (IC50) of ssDNA 5’-GTGGG-3’ was calculated as 2754.00 ± 682.00 nM, confirming its competitive role in disrupting SCAF4-RNAP2 interaction (Fig. 1i; Supplementary Fig. 1e). As a result of its potent inhibitory efficacy and robust binding affinity, ssDNA 5’-GTGGG-3’ (hereinafter referred to as the SCAF4 lead sequence (SCAF4_LS)), was selected as the primary iAptamer in this study.

### Specific interaction of SCAF4_LS binding to SCAF4

To gain insight into the interaction between SCAF4_LS and SCAF4, a molecular dynamics (MD) simulation was carried on the initial complex model generated from docking. The production phase simulations were performed for a total time of 500 ns at a constant temperature of 298 K. The binding position of SCAF4_LS on SCAF4 shifted, but still stayed tightly associated with SCAF4 along the simulation and final entrance to a stable binding pose after 300 ns MD simulation. The binding poses of SCAF4_LS in the last 100 ns (400-500 ns) were summarized in Figure 2a. The obtained root mean square fluctuation (RMSF) values of each nucleotide from all calculated conformations of SCAF4_LS were plotted against the simulation time from 400 ns to 500 ns for the analysis of nucleotide stability (Fig. 2b). The trajectory clearly shows that the 3’ terminus of the DNA molecule exhibited greater flexibility compared to its 5’ terminus, while G1, T2, and G3 bases at the 5’ terminus exhibited stable binding with SCAF4 protein, as evidenced by the RMSF values, which were used to measure the average deviations of each nucleotide from the averaged structure in 400-500 ns (Fig. 2b). This observation was further corroborated by plotting the RMSD values of individual nucleotides against the simulation time between 400 to 500 ns (Fig. 2c). Collectively, our simulation results suggest that SCAF4_LS binds to SCAF4 protein in a stable manner and that the first three nucleotides of the aptamer exhibit a particularly heightened interaction with the protein, while the last two nucleotides slide over the protein surface (Fig. 2a).

**Figure 2:**
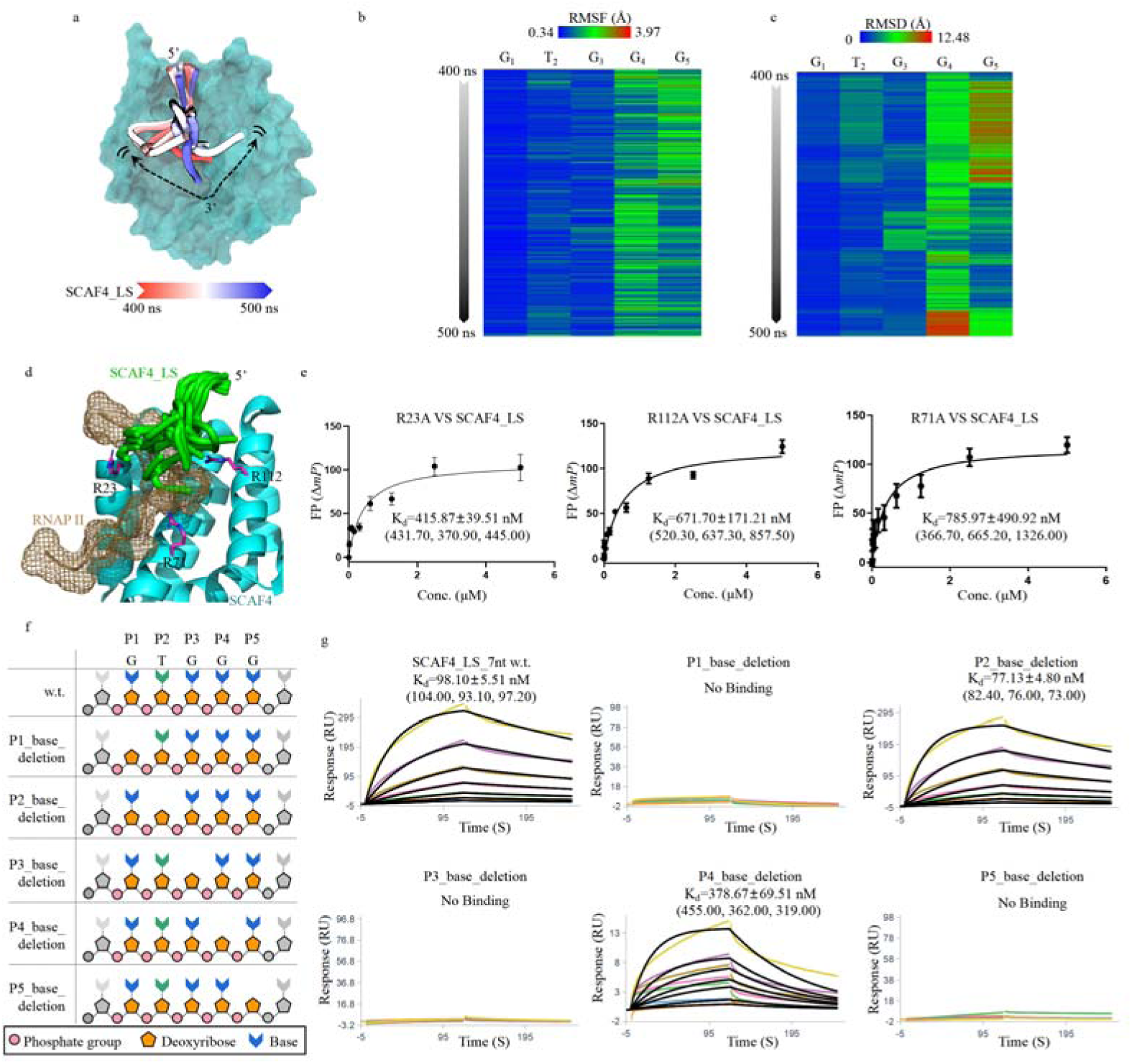
SCAF4_LS binding at the interface between SCAF4 and RNAP2. **a**) Molecular simulation of SCAF4_LS binding to SCAF4 protein. The colors of SCAF4_LS are color-coded based on the time interval from 400 ns to 500 ns. **b**) Trajectory plot showing the RSMF of five nucleotides over the 400-500 ns time scale. **c**) Trajectory plot showing the RSMD of five nucleotides over the 400-500 ns time scale. **d**) Overlay of SCAF4_LS sequence and RNAP2 on the surface of SCAF4 protein. Conformations of SCAF4_LS are represented in green cartoon, while RNAP2 peptide is depicted in sand-colored mesh. **e**) Fluorescence polarization measurements where 50 nM FAM-labeled SCAF4_LS was titrated with increasing amounts of SCAF4 mutants. Affinities were determined as 415.87 ± 39.51 nM, 671.70 ± 171.21 nM and 785.97 ± 490.92 nM for R21A, R112A and R71A respectively. All experiments were performed with three biological replicates. Affinities of each replicate were indicated in the bracket. The non-linear fit model and the one-site specific binding to fluorescent population ratio were used for these calculations. **f**) Schematic diagram of SCAF4_LS with deletion of base groups at each position. **g**) SPR results showing binding of wild type and base group-deleted SCAF4_LS to SCAF4. Average affinities were generated from three independent experiments, with affinities of each experiment indicated in the bracket.

SCAF4_LS has a marked predilection for engaging with a positively charged groove located on the PPI interface of SCAF4, as demonstrated in the molecular simulation assay (Fig. 2d). Superposition of the simulated SCAF4_LS aptamer-SCAF4 model and RNAP2-SCAF4 complex structure (6XKB) reveals a clear overlap between the RNAP2-CTD and SCAF4_LS aptamer (Fig. 2d). This supports the competitive role of SCAF4_LS in blocking the interaction between SCAF4 and RNAP2 (Fig. 2d). To determine the binding specificity of SCAF4_LS sequence to the PPI interface, critical residues on the SCAF4 interface were substituted, and their binding to SCAF4_LS was characterized using a fluorescence polarization assay. Three arginine residues (R23, R71, and R112), which are known to be involved in SCAF4 recognition of RNAP2^53^, were substituted with alanines, resulting in weaker interaction with SCAF4_LS and reducing the binding affinity by over two-fold (Fig. 2e; Supplementary Fig. 1f-h).

In parallel, to investigate the base specificity SCAF4_LS-SCAF4 protein interaction, a series of base-group deletions for each individual nucleotide within the SCAF4_LS sequence was synthesized (Fig. 2f). These deletions were incorporated into a seven-based backbone sequence (5’-GGTGGGG-3’) containing two additional nucleotides, required in DNA synthesis. The affinity of each base deletion of SCAF4_LS sequence to SCAF4 protein was then assessed by SPR (Fig. 2g; Supplementary Fig. 2a-f). Significantly, the elimination of guanine groups from nucleotides G1, G3, and G5 drastically abrogated interaction between SCAF4_LS and SCAF4, underscoring the critical roles of these three guanine base groups. Additionally, removing the guanine group from G4 led to reduced affinity. However, SCAF4_LS lacking the thymine group at T2 still maintained its affinity to SCAF4. These findings collectively demonstrate the sequence-specific binding of SCAF4_LS aptamer to SCAF4 protein. Moreover, our results indicate that the SCAF4_LS sequence and the phosphorylated RNAP2 peptide share a common binding region on SCAF4, supporting the competitive nature of the SCAF4_LS aptamer in impeding RNAP2-SCAF4 interactions.

### Optimization of SCAF4_LS

Having identified the initial SCAF4_LS aptamer with binding affinity and inhibitory effects, further optimization was performed by expanding the sequence to generate 64 (4^3^) additional sequences, each with three nucleotides added to the termini (Fig. 3a). These expanded sequences were then evaluated for their inhibitory effects using a competitive fluorescence polarization assay with the FITC-labeled CTD peptide acting as the fluorescence probe and the extended sequences as competitors (Fig. 3b). Among these expanded sequences, three sequences (B2_8nt, D1_8nt, and F1_8nt) displayed significant signal reductions, indicating their pronounced competitiveness in blocking SCAF4-RNAP2 interactions (Fig. 3c). The inhibitory effects of B2_8nt, D1_8nt, and F1_8nt on SCAF4-RNAP2 interactions were further determined as 1277.50 ± 742.05 nM, 1265.67 ± 864.02 nM and 553.73 ± 106.37 nM, respectively, by competitive FP assays (Supplementary Fig. 3a). Concurrently, the interactions of these 64 expanded sequences with SCAF4 proteins were investigated using SPR. It was found that multiple sequences, including B2_8nt, D1_8nt, and F1_8nt, showed significant binding to SCAF4 with response units (RU) over 200 (Fig. 3d). The binding affinities of F1_8nt and B2_8nt to SCAF4 protein were further quantified as 88.60 ± 15.09 nM and 296.67 ± 48.13 nM, respectively, by SPR (Supplementary Fig. 3b, c). Based on these findings, two sequences, F1_8nt and B2_8nt, were selected as the seed sequences in parallel for the second round of aptamer optimization owing to their significant binding and inhibitory abilities on SCAF4-RNAP2 interactions (Fig. 3e).

**Figure 3:**
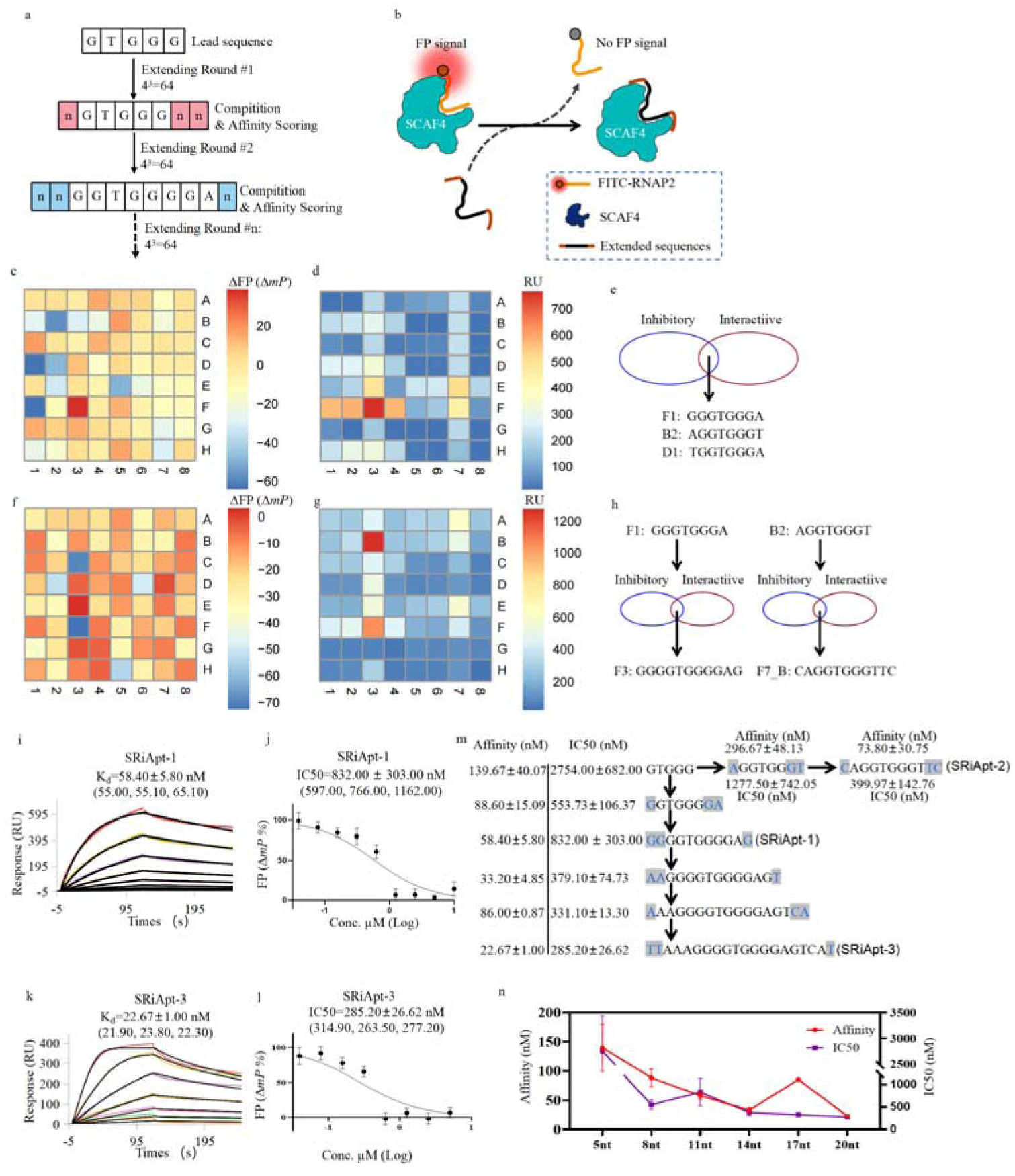
Optimization of SCAF4_LS to improve its affinity and inhibitory capacity. **a**) Workflow illustrating the sequence optimization procedures through the addition of supporting nucleotides to SCAF4_LS. **b**) Schematic representation of the competition assay used for the selection of extended sequences. **c**) Evaluation of the inhibitory efficacy of the expanded 64 sequences in interfering with SCAF4 and RNAP2 interaction, utilizing a competitive FP assay. All 64 DNA sequences were tested at a final concentration of 1.5 µM. **d**) Direct interactions of the expanded 64 sequences with SCAF4 assessed by SPR spectroscopy. Proteins were exposed to 10 µM of the 30 top-scoring 5-nt sequences. **e**) Selection of DNA sequences exhibiting both binding affinity and inhibitory effects. **f**) Assessment of the inhibitory effects of the second round of expanded 64 sequences on the SCAF4 and RNAP2 interaction. **g**) Direct interactions of the second round of expanded 64 sequences to SCAF4. **h**) Selection of DNA sequences demonstrating both binding affinity and inhibitory effects. **i)** Standard kinetics SPR assay demonstrating the binding of SCAF4_F3_11nt (SRiApt-1) to immobilized SCAF4. This experiment was conducted with three replicates. Average affinity value was 58.40 ± 5.80 nM. Affinities for each replicate were indicated in the bracket. **j)** Dose-response curves depicting the inhibition of RNAP2 interaction with SCAF4 by SCAF4_F3_11nt (SRiApt-1). Half-maximal inhibitory concentrations were determined from three independent experiments, each containing three independent replicates. IC50 values were calculated using GraphPad Prism software (version 7) with the Log(inhibitor) vs. normalized response model. IC50 values of all experiments are indicated in the bracket (n=3, mean ± SD). **k)** Standard kinetics SPR assay demonstrating the binding of SRiApt-3 to immobilized SCAF4. This experiment was conducted with three replicates. Average affinity value was 22.67 ± 1.00 nM. Affinities for each replicate were indicated in the bracket. **l)** Dose-response curves depicting the inhibition of RNAP2 interaction with SCAF4 by SRiApt-3. Half-maximal inhibitory concentrations were determined from three independent experiments, each containing three independent replicates. **m**) Summary of aptamers optimized from the lead sequence. Both affinities and IC50s of selected aptamers were labeled on the left, and the sequence information was indicated on the right. New residues iterated from the previous sequence were labeled in gray. **n**) Iterations of affinities and IC50s for SRiApt-3, with affinity shown in red and IC50 displayed in blue.

In the subsequent round of aptamer refinement originating from F1_8nt, two remarkably competitive aptamer sequences (C3_11nt and F3_11nt) were identified through a competitive FP assay, mirroring the prior optimization process (Fig. 3f). Concurrently, F3_11nt exhibited the most substantial binding to the SCAF4 protein in the SPR screening assay (Fig. 3g). Consequently, the F3_11nt sequence was selected for further evaluation (Fig. 3h). The binding affinity of F3_11nt to SCAF4 was determined to be 58.40 ± 5.80 nM (Fig. 3i; Supplementary Fig. 3d). The inhibitory effect of SCAF4_F3_11nt aptamer on SCAF4-RNAP2 interactions was verified through competitive FP assay, showing an IC50 value of 832.00 ± 303.00 nM and thus showcasing its enhanced inhibitory potential compared to the parent SCAF4_LS aptamer (Fig. 3j; Supplementary Fig. 3e). Notably, despite the guanine-rich nature of F3_11nt, the investigation of G-quadruplex structures across various K^+^ concentrations did not yield any observed G-quadruplex structures within this sequence (Supplementary Fig. 3f). The binding specificity of the SCAF4_F3_11nt aptamer to SCAF4 protein was then explored by examining its interactions with three unrelated proteins (streptavidin, BSA, and IgG), revealing no observable interactions and, thus, affirming its elevated binding specificity (Supplementary Fig. 3g).

Simultaneously, extended sequences derived from B2_8nt underwent screening via competitive FP assay. Many sequences demonstrated substantial reductions in FP signal, leading to the selection of three sequences with the most pronounced reductions (F7_11nt_B, H7_11nt_B, and G8_11nt_B) for further analysis. The inhibitory potency of each sequence towards SCAF4-RNAP2 interaction was determined to be 399.97 ± 142.76 nM, 437.50 ± 129.49 nM, and 459.57 ± 188.98 nM, respectively, through competitive FP assays (Supplementary Fig. 4a-c). Moreover, the binding affinity between each sequence, F7_11nt_B, H7_11nt_B, and G8_11nt_B, and SCAF4 protein was quantified via SPR as 73.80 ± 30.75 nM, 62.03 ± 14.32 nM, and 57.03 ± 1.65 nM, respectively (Supplementary Fig. 4d-f). Owing to the substantial affinity of each sequence to SCAF4 and potent inhibitory effect of each on SCAF4-RNAP2 interactions, SCAF4_F3_11nt and SCAF4_F7_11nt_B aptamers were designated as SCAF4-RNAP2 inhibitory aptamer 1 (SRiApt-1) and SriApt-2, respectively (Fig. 3h, m).

At last, SRiApt-1 underwent further optimization to maximize its binding and inhibitory capabilities. Through additional three rounds of iterative refinement, we successfully identified a 20-base DNA sequence with an affinity of 22.67 ± 1.00 nM to SCAF4 and an IC50 of 285.20 ± 26.62 nM for blocking SCAF4-RNAP2 interactions (Fig. 3k,l; Supplementary Fig. 5). Consequently, this aptamer was designated as SCAF4-RNAP2 inhibitory aptamer 3 (SRiApt-3).

In summary, employing the Blocker-SELEX pipeline and implementing a series of iterative optimizations on the lead sequence, we achieved a continuous enhancement in both the affinity and inhibitory capacity of the aptamers (Fig. 3m). While there were occasional fluctuations in the trends of affinity or inhibitory, the overall effectiveness of the iterative optimization approach to improve the performance of aptamers was robustly validated (Fig. 3n).

### SCAF4_LS sequence was recaptured in developing aptamers for SCAF8

SCAF8, a crucial paralog of SCAF4, shares a comparable CID domain with an overall RSMD of 0.266 Å and a PPI interface for RNAP2 recognition^53,54^ (Supplementary Fig. 6a). Concomitant knockout of human SCAF4 and SCAF8 results in altered poly-A selection and subsequent early termination, leading to expression of truncated mRNAs and proteins lacking functional domains, ultimately causing cell death^52^. In light of these observations, we embarked on selecting aptamer inhibitors capable of blocking SCAF8-RNAP2 interaction by again using the Blocker-SELEX pipeline. Interface details were obtained from the SCAF8-RNAP2 complex structure (PDB code: 3D9M). The binding potential of 5-nt ssDNA sequences was scored by Glide docking. As anticipated, the structural similarity between SCAF4-CID and SCAF8-CID caused the SCAF4_LS sequence to emerge as one of the top-scoring sequences (Fig. 4a).

**Figure 4:**
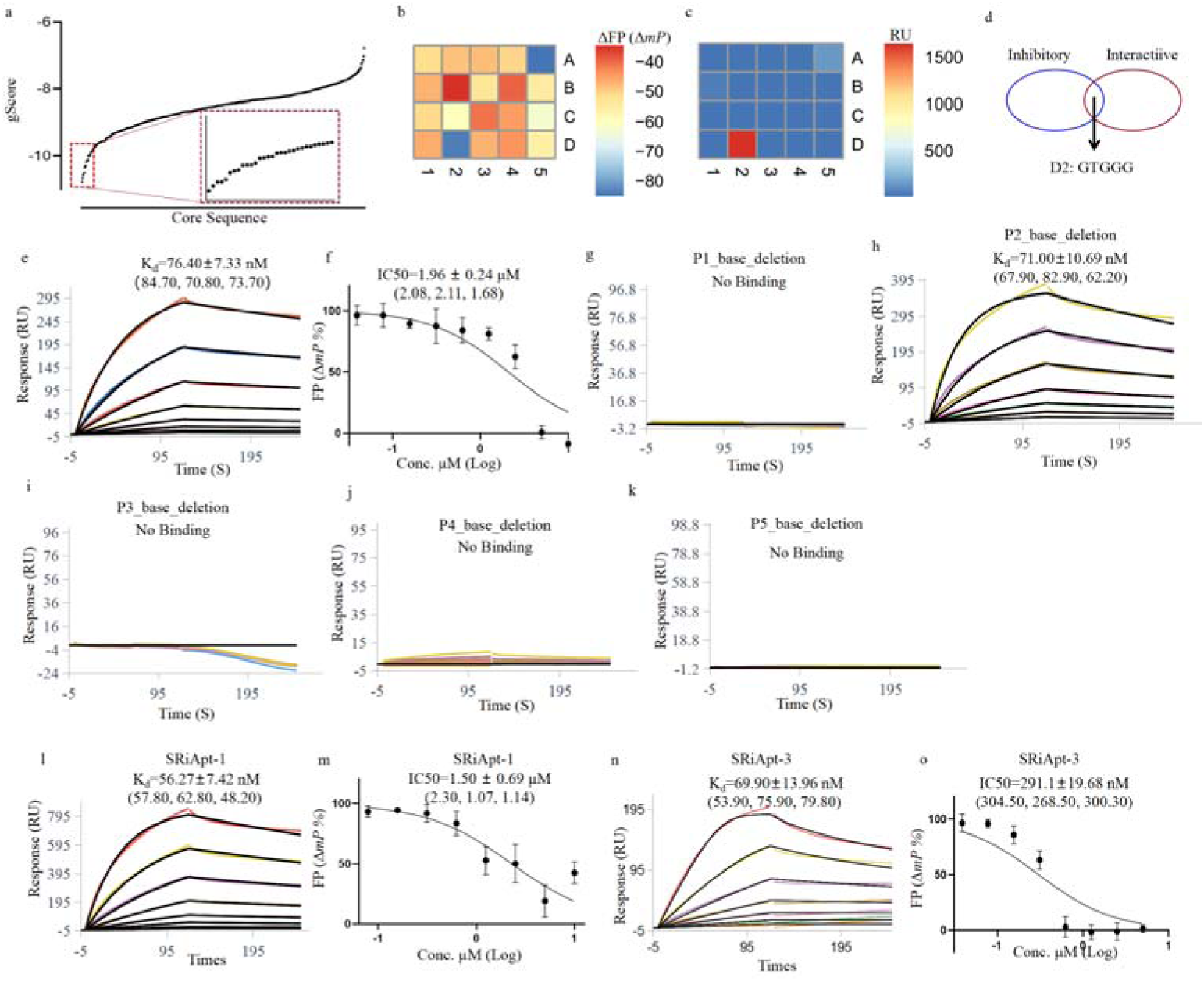
Design of aptamers for blocking RNAP2 and SCAF8 interactions. **a**) Docking scores of 5-nt sequences to SCAF8 generated by the Glide pipeline. The top 20 sequences were enlarged in the inserted panel. **b**) Assessment of the inhibitory effects of the top 20 sequences on the interaction between SCAF8 and RNAP2 via competitive fluorescent polarization assay. **c**) Evaluation of the direct interactions between the top 20 sequences and SCAF8 using SPR spectroscopy. Proteins were exposed to 10 μM of the top 20 highest scoring 5-nt sequences. RU, response units. **d**) Selection of DNA sequences exhibiting both binding affinity and inhibitory effects. **e**) Standard kinetics SPR assay demonstrating the binding of SCAF4_LS (D2_5nt) to immobilized SCAF. This experiment was performed with three replicates. Average affinity value was 76.40 ± 7.33 nM. Affinities of each replicate were indicated in the bracket. **f**) Dose-response curves of SCAF4_LS (D2_5nt) depicting the inhibition of RNAP2 interaction with SCAF8. The averaged IC50 value was calculated from all experiments indicated in brackets (n=3, mean ± SD). **g-k**) SPR results illustrating the binding of base group-deleted SCAF4_LS (D2_5nt) sequences to SCAF8. Average affinities were generated from three independent experiments, with affinities of each experiment indicated in the bracket. **l**) SPR results demonstrating the binding of SCAF4_F3_11nt to SCAF8. The average affinity was 58.40 ± 5.80 nM from three independent experiments. Affinities for each replicate are indicated in the bracket. **m**) Dose-response curves of SRiApt-1 illustrating the inhibition of RNAP2 interaction with SCAF8. The half-maximal inhibitory concentrations were determined from three independent experiments indicated in the bracket (n=3, mean ± SD). **n**) SPR results demonstrating the binding of SRiApt-3 to SCAF8. The average affinity was 69.90 ± 13.96 nM from three independent experiments. Affinities for each replicate are indicated in the bracket. **o**) The inhibition of RNAP2 interaction with SCAF8 by SRiApt-3 with an IC50 of 291.10 ± 19.68 nM.

In order to evaluate the inhibitory efficacy of these top-scoring sequences, SCAF8-RNAP2 interaction was first recapitulated using a fluorescence polarization assay, yielding an affinity of 622.40 nM (Supplementary Fig. 6b). For competition screening, FP signals (> 60 Δ mP) were generated by combining 2 µM SCAF8-CID with 40 nM FITC-labeled CTD peptide solutions. The top 20 sequences were individually screened for their inhibitory efficacy by adding 10 µM of each sequence into the FP solutions. Among the top 20 sequences, D2_5nt (SCAF4_LS) and A5_5nt sequences exhibited the most potent competition with SCAF8-RNAP2 interaction, as determined by the competitive fluorescence polarization assay (Fig. 4b). The binding between each of 20 sequences to SCAF8 protein was further assessed using SPR, revealing that D2_5nt (SCAF4_LS) bound to SCAF8 protein with RU over 1000, while the remaining sequences displayed no significant responses (Fig. 4c). Furthermore, the affinity between D2_5nt (SCAF4_LS) and SCAF8 was quantified as 76.40 ± 7.33 nM, using SPR (Fig. 4e; Supplementary Fig. 6c). The inhibitory effect of D2_5nt (SCAF4_LS) on SCAF4-RNAP2 interaction was determined to have an IC50 value of 1960.00 ± 240.00 nM (Fig. 4f; Supplementary Fig. 6d). Collectively, D2_5nt (SCAF4_LS), with both significant binding affinity and inhibitory effects, was selected as the lead sequence for targeting SCAF8-RNAP2 interaction (Fig. 4d).

To assess the contribution of each nucleotide base of D2_5nt (SCAF4_LS) to SCAF8 binding, the affinity of each base group-deleted variant of D2_5nt (SCAF4_LS) was investigated (Fig. 2f; Fig. 4g-k; Supplementary Fig. 7). Similar to the interaction between SCAF4_LS and SCAF4, the removal of guanine groups at positions P1, P3, P4, and P5 resulted in substantial reductions in affinity, indicating the critical role of these four guanine base groups in D2_5nt (SCAF4_LS) recognition. We further examined the binding affinities of optimized SRiApt-1/2/3 to SCAF8, and their affinities were determined as 56.27 ± 7.42 nM, 90.50 ± 25.80 nM and 69.90 ± 13.96 nM, respectively (Fig. 4l, n; Supplementary Fig. 6e-g). The IC50s of SRiApt-1/2/3 aptamers in blocking SCAF8-RNAP2 interaction were determined as 1.50 ± 0.69 μM, 2.69 ± 1.43 μM and 291.1 ± 19.68 nM respectively (Fig. 4m, o; Supplementary Fig. 6h-j). These findings support that SRiApt-1/2/3 aptamers simultaneously block SCAF4 or SCAF8-RNAP2-CTD interactions.

Ultimately, our study highlights the reproducibility of the Blocker-SELEX approach in generating iAptamers that specifically disrupt the interactions between SCAF4/SCAF8 and RNAP2-CTD.

### Binding of SRiApt-1/3 aptamers to SCAF4/SCAF8 at the cellular level

To investigate the interactions between the selected inhibitory aptamers and SCAF4/SCAF8 proteins within a cellular context, an immunoprecipitation (IP) assay was performed using SRiApt-1/3 aptamers. Streptavidin-coated magnetic beads were separately immobilized with biotinylated SRiApt-1/3 aptamers and the reverse complementary sequences (5’-CTCCCCACCCC-3’ or 5’-ATGACTCCCCACCCCTTTAA-3’) as the control aptamers. Subsequently, the aptamer-loaded beads were incubated with HCT116 cell lysates for 30 minutes, followed by three rounds of washing with PBS buffer to remove nonspecifically bound components. The proteins that specifically bound to the aptamer-loaded beads were then detected via SDS-PAGE and Western blot analysis using specific antibodies targeting SCAF4 or SCAF8. The IP assays demonstrated that the SRiApt-1 or SRiApt-3 aptamer-loaded beads successfully captured SCAF4 or SCAF8 proteins, while the control sequences showed no significant protein enrichment, indicating direct interaction between SRiApt-1/3 aptamers and these proteins within the cellular context (Fig. 5a). As SCAF4/8-RNAP2 interactions predominantly occur within the nucleus, it was crucial to evaluate the nuclear accumulation of the selected inhibitory aptamers. Hence, fluorescence localization assays were employed to examine the subcellular distributions of SRiApt-1/3 aptamers. FAM-labeled SRiApt-1/3 aptamers were introduced into living cells through SM-102, and their distribution was monitored over time. Within 6 hours of transfection, the FAM-labeled SRiApt-1/3 aptamers exhibited significant accumulation in the nucleus, as indicated by strong green fluorescence signals. This observation confirms the rapid nuclear localization of the SRiApt-1/3 aptamers (Fig. 5c; Supplementary Fig. 8a). Furthermore, to investigate whether the SRiApt-1/3 aptamers could disrupt SCAF4/8-RNAP2 interaction in a cellular context, a CoIP assay was performed, using SCAF4 as an example. HA-tagged SCAF4-CID was exogenously overexpressed in HEK293 cells, and endogenous RNAP2 protein was co-precipitated using an anti-HA antibody, but the co-precipitation was significantly reduced upon treatment with the SRiApt-1/3 aptamers. These results conclusively validate the inhibitory potential of SRiApt-1/3 aptamers in blocking SCAF4-RNAP2 interaction (Fig. 5b).

**Figure 5:**
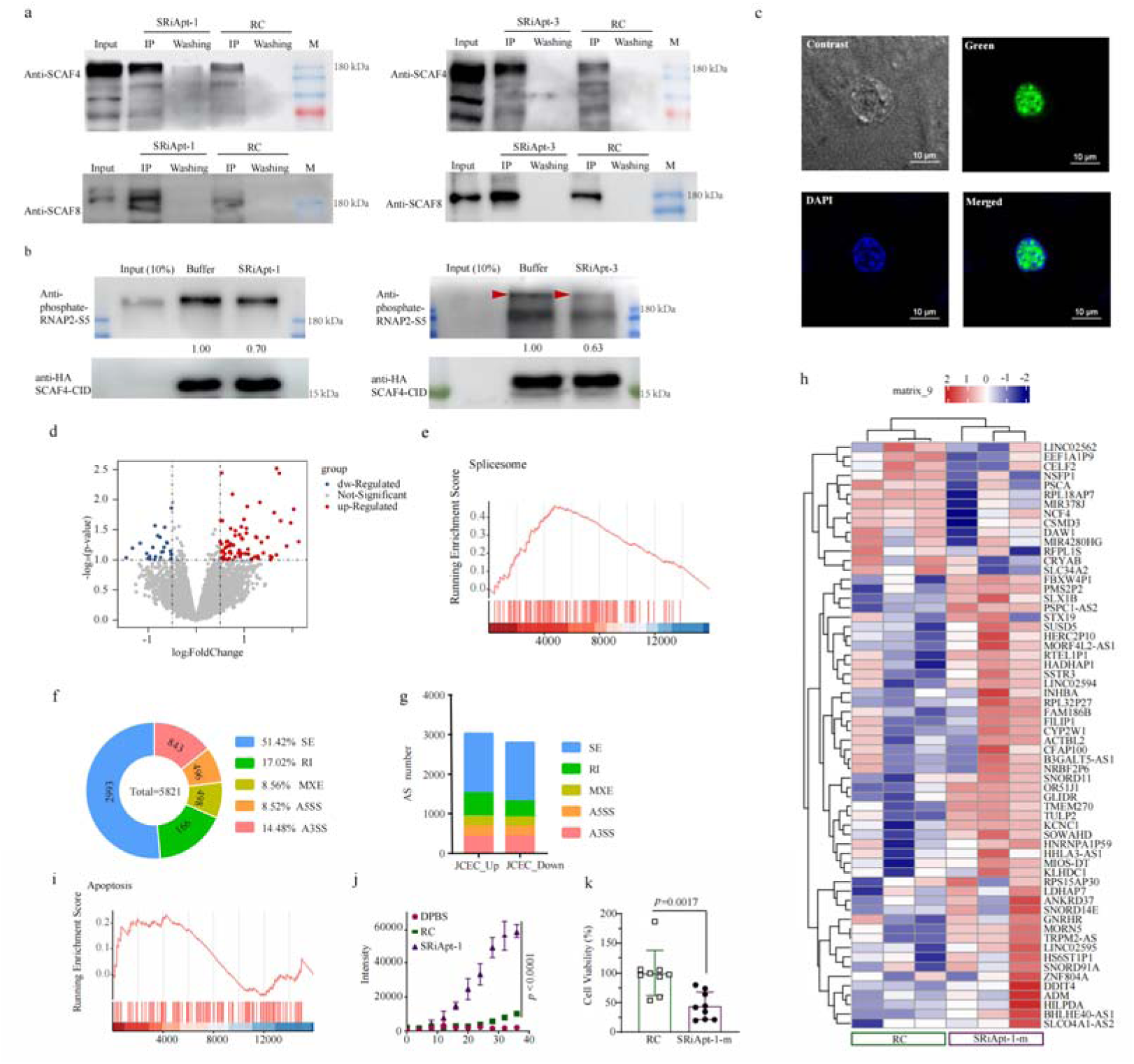
Biological characterizations of the SRiApt-1/3 aptamers. **a**) Co-IP assays using Biotin-SRiApt-1/3 aptamers or the corresponding Biotin-RC sequences. **b**) CoIP assays revealing the inhibitory role of SRiApt-1/3 aptamers in blocking the interaction between SCAF4 and RNAP2. **c**) Intracellular localization of FAM-labeled SRiApt-1 aptamer following transfection with LNPs. Aptamers are highlighted in green, while DAPI staining is shown in blue. Scale bar: 10 μm. **d**) A volcano plot representing the differential expression of genes between the treated group with SRiApt-1 aptamers to that in the control group treated with the RC sequence, based on their statistical significance and fold change. Genes that exhibited significant changes were selected using log_2_FoldChange criteria (>0.5 or < −0.5) and a p-value threshold (< 0.1). Each dot on the plot represents an individual gene, and their color indicates the corresponding regulation as specified in the legend. The log_2_FoldChange was calculated and transformed from the ratio of gene expression in the treated group with SRiApt-1 aptamers to that in the control group treated with the RC sequence. **e**) Gene Set Enrichment Analysis (GSEA) for KEGG enrichment (NES=1.665, p value <0.05) of the Spliceosome pathway after treatment with SRiApt-1 aptamers. Data analyzed by one-tailed Fisher’s exact test. **f**) Pie plot displaying the number and ratio of ASEs belonging to each of the main alternative splicing categories. **g**) Bar plot showing the number and ratio of up-/down-regulated ASEs belonging to each of the main alternative splicing categories. The color represents various alternative splicing (AS) categories. The horizontal axis represents upregulation or downregulation of AS events, and the vertical axis represents the number of alternative splicing events for each type. JCEC represents AS event detection using both Junction Counts and reads on target. **h**) Heatmap representing significant upregulated and downregulated mRNA or non-coding RNA with SRiApt-1 aptamers treatment (absolute value of Log_2_ fold change >1), compared to the RC sequence treating. **i**) GSEA for KEGG gene sets associated with the Apoptosis pathway (NES=0.83, p value <0.05) after treated with SRiApt-1 aptamers. Data were analyzed by one-tailed Fisher’s exact test. **j**) Representative apoptotic curves of HCT116 cells treated with SRiApt-1 or RC control for 3 days under Annexin V staining. n=3, mean ± SD. Two-sided Student’s t-test. **k**) Cell viability assessed using the CCK8 assay when cells were treated with SRiApt-1-m or RC aptamers at a working concentration of 1 µM. n=3, mean ± SD. Two-sided Student’s t-test.

### SRiApt-1 aptamer altered gene transcription and RNA splicing

Previous studies have demonstrated that simultaneous depletion of SCAF4 and SCAF8 disrupts the gene transcriptome and promotes alternative splicing events^52^. To validate the targeted effect of those selected inhibitory aptamers on gene transcription and mRNA splicing by inhibiting the interaction between RNAP2 and SCAF4/SCAF8 proteins at the cellular level, we employed the SRiApt-1 aptamer. Initially, we enhanced the stability of SRiApt-1 by introducing two 2’-O-Me modifications, resulting in SRiApt-1-m (5’-G(2’-O-Me)GGGTGGGGA(2’-O-Me)G-3’) (Supplementary Fig. 3f). Subsequently mRNA sequencing was conducted on HCT116 cells treated with SRiApt-1-m, or the reverse complementary sequence, for 36 hours. Analysis of RNA-seq data revealed significant alterations in the transcriptome of HCT116 cells upon SRiApt-1-m aptamer treatment (Fig. 5d). Specifically, 97 genes were identified as differentially expressed with 72 genes upregulated and 25 genes downregulated (P value < 0.1, Fig. 5d). Gene set enrichment analysis (GSEA) was also performed, showing upregulation of the spliceosome pathway upon SRiApt-1-m treatment (Fig. 5e). Further examination of alternative splicing events based on the RNA-seq data revealed a substantial increase, totaling 5821 events, with over 50% classified as skipped exons (SE), subsequent to SRiApt-1-m aptamer treatment (Fig. 5f, g). Additionally, numerous non-coding RNAs (ncRNAs) were identified in the RNA-seq data when aligned with the human genome (Fig. 5h). These findings align with a recent study that demonstrated significant alternative splicing events resulting from the double knockout of SCAF4 and SCAF8^52^.

Since one of the crucial cellular processes influenced by alternative splicing is programmed cell death, or apoptosis, we investigated the RNA-seq data and observed activation of the apoptosis signaling pathway following treatment with the SRiApt-1-m aptamer (Fig. 5i). To validate this observation, a cell apoptosis assay was performed using HCT116 cells. Cells were exposed to SRiApt-1-m, or the reverse complementary sequence, using vehicles at a final concentration of 1 µM. Subsequently, apoptotic cells were labeled with Annexin V, a marker of apoptosis. A noticeable increase in apoptotic cells was observed compared to the control sequence, confirming the ability of the SRiApt-1-m aptamer to promote apoptosis in HCT116 cells (Fig. 5j, Supplementary Fig. 8d). Furthermore, the CCK8 assay provided additional confirmation of the inhibitory effects of SRiApt-1 or SRiApt-1-m aptamers on HCT116 cell proliferation (Fig. 5k, Supplementary Fig. 8e). Collectively, these findings substantiate the potential therapeutic utility of the SRiApt-1-m aptamer inhibitor in impeding tumor growth. Consequently, these observations support the on-target effect of SRiApt-1 and establish its significance as a valuable chemical tool for investigating RNAP2-SCAF4/SCAF8 interactions.

### Blocker-SELEX pipeline is feasible against WDR5-MYC interaction

To assess the versatility of Blocker-SELEX in selecting aptamers targeting TF interactions, we aimed to develop inhibitory aptamers capable of blocking the interactions of the intrinsically disordered oncogenic MYC protein. Among the cofactors known to regulate MYC, WD40-repeat protein 5 (WDR5) stands out as a key determinant for recruiting MYC to chromatin, a process essential for MYC’s oncogenic properties^55^. Disrupting this interaction has been shown to impede MYC binding at approximately 80% of its chromosomal locations, leading to the deactivation of its oncogenic properties^55,56^. Utilizing the Blocker-SELEX pipeline, we first prepared the WDR5-MYC binding interface from the crystal structure of WDR5-MYC complex (PDB code: 4Y7R) (Supplementary Fig. 9a). Then we calculated the binding potentials of ssDNA sequences to the interface of WDR5. The top 40 sequences were assessed for their competition to WDR5-MYC interactions by a competitive FP assay (Supplementary Fig. 9b). In this assay, we utilized an N-terminal FITC-labeled MYC peptide, along with purified WDR5 proteins, to establish the initial FP signal (Supplementary Fig. 9c). Subsequently, each of the synthesized 40 sequences was introduced individually to the WDR5-MYC mixture at a concentration of 10 µM. Among those sequences, 5’-GGACC-3’ exhibited a modest, yet significant, reduction in the FP signal and was consequently selected as the lead sequence (Fig. 6a). The inhibitory effect of the lead’s bivalent form, 5’-GGACCGGACC-3’, was further quantified, resulting in an IC50 value of 8.50 ± 4.25 µM, indicative of roughly two-thirds reduction in the FP signal at 10 µM (Fig. 6b; Supplementary Fig. 9d). The sequences underwent further optimization by incorporating additional supporting bases to the lead sequence in accordance with our standard protocol (Supplementary Fig. 9e, f, g). Through five rounds of iteration using the competitive FP assay, a stem-loop folded sequence, 5’-GGGGGCTGGACCCCTCAACT-3’, was identified (Fig. 6c). Secondary structure analysis indicated a stem-loop folding with a calculated ΔG of −4.30 kcal/mol. This sequence, designated as WDR5-MYC-iAptamer-1 (WMiApt-1), exhibited a substantially lower IC50 value of 431.30 ± 133.61 nM in disrupting the WDR5-MYC interaction (Fig. 6d; Supplementary Fig. 9f). Then, we tried to improve the inhibitory capacity by stabilizing the stem region through the design of four variants with improved ΔG values, yet all exhibited markedly weaker inhibitory capacity, suggesting the predicted stem could not accommodate change (Fig. 6e; Supplementary Fig. 9h). The affinity between WMiApt-1 and WDR5 was determined using the FP assay, yielding a value of 797.37 ± 95.16 nM (Fig. 6f; Supplementary Fig. 9i). This binding was also validated through SPR with a value of 138.20 ± 86.73 nM (Fig. 6g; Supplementary Fig. 9j).

**Figure 6:**
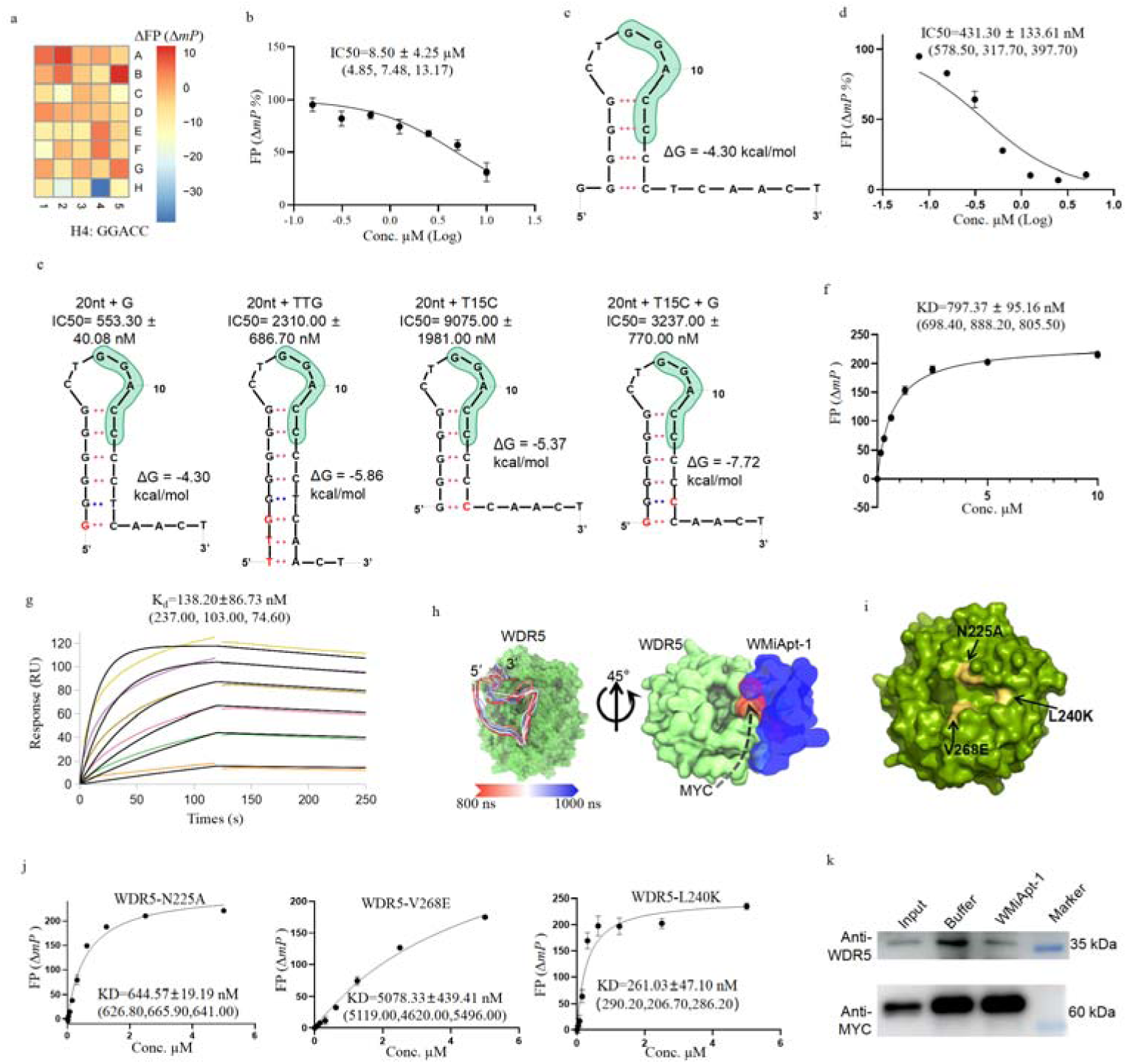
Design of inhibitory aptamers targeting the oncogenic WDR5-MYC interaction. **a**) Assessment of the inhibitory effects of the top 40 sequences on the interaction between WDR5 and MYC via competitive fluorescent polarization assay. **b**) Dose-response curves depicting the inhibition of the interaction between WDR5 and MYC by the sequence “5’-GGACCGGACC-3’”. Half-maximal inhibitory concentrations were determined from three independent experiments, each containing three independent replicates. IC50 values were calculated using GraphPad Prism software (version 7) with the Log(inhibitor) vs. normalized response model. IC50 values of all experiments are indicated in the bracket (n=3, mean ± SD). **c**) The secondary structure of WMiApt-1 generated by the DNA folding form with a ΔG of −4.30 kcal/mol and melting temperature of 62.8 ℃. The lead sequence was highlighted in green. **d**) Inhibition of WMiApt-1 to MYC-WDR5 interaction. The half-maximal inhibitory concentrations were determined from three independent experiments, each containing three independent replicates. IC50 values of all experiments are indicated in the bracket (n=3, mean ± SD). **e**) Summary of IC50s of WMiApt-1 variants in blocking WDR5-MYC interactions. The Mutated bases were labeled in red. IC50 values were generated from three independent replicates. **f**) Fluorescence polarization measurements of the interaction between WDR5 and WMiApt-1. The affinity was determined as 797.37 ± 95.16 nM using the FP (ΔmP) signals with three biological replicates. Affinities of each replicate are indicated in the bracket. **g**) Standard kinetics SPR assay demonstrating the binding of WMiApt-1 to immobilized WDR5. This experiment was performed with three replicates. Average affinity value was 138.20 ± 86.73 nM. Affinities of each replicate were indicated in the bracket. **h**) A model of WDR5-WMiApt-1 complex generated by molecular simulation with the colors of WMiApt-1 color-coded based on the time interval from 800 ns to 1000 ns (left). Its overlap with MYC was shown on the right with WDR5 in limon, WMiApt-1 in blue, and MYC in red. **i, j**) The locations of three key residues critical for WDR5-MYC interactions were shown in yellow. Fluorescence polarization measurements of the interaction between WMiApt-1 and WDR5 proteins with the three key mutants. **k**) CoIP assays revealing the inhibitory role of WMiApt-1 in blocking the interaction between WDR5 and MYC.

Furthermore, a comprehensive model of the WDR5 and WMiApt-1 complex was generated through MD simulations, supporting the competitive role of WMiApt-1 in the WDR5-MYC binding (Fig. 6h). To validate this simulation model, three mutations previously identified as crucial to the WDR5-MYC interaction were introduced to WDR5 proteins^57^(Fig. 6i). The pivotal roles of these three residues in recognizing MYC were reaffirmed through FP assays; all three mutations attenuated the affinity between WDR5 and MYC (Fig. 6i; Supplementary Fig. 9l-n). In comparison to the wild-type WDR5, which exhibited an affinity of 430.00 ± 130.00 nM to WMiApt-1 (Supplementary Fig. 9c), two of the three residues (N225 and V268), showing reduced affinities to WMiApt-1, emerged as pivotal for WMiApt-1 binding (Fig. 6i; Supplementary Fig. 9o-q). This validation supports the simulation model, confirming the binding surface of WMiApt-1 on the WDR5 interface. Although the L240K mutation improved the affinity, it is reasonable to posit that despite an overlap between WMiApt-1 and MYC on the WDR5 interface, distinctions in the recognition patterns between WMiApt-1 and WDR5 persist. Subsequently, we explored the binding specificity of WMiApt-1 to WDR5 protein by investigating its interactions with two unrelated proteins, streptavidin and IgG. This analysis revealed no specific interactions with these unrelated proteins, underscoring WMiApt-1’s binding specificity to WDR5 (Supplementary Fig. 9k). These findings collectively highlight the potential of WMiApt-1 aptamer in disrupting WDR5-MYC interaction at the cellular level. To experimentally validate the inhibitory effects of WMiApt-1 on WDR5-MYC interaction within cells, CoIP studies were conducted. By exogenous over-expression of MYC protein in HEK293 cells, the WDR5-MYC complex was captured by anti-MYC antibody, but disrupted upon WMiApt-1 aptamer treatment (Fig. 6j; Supplementary Fig. 9r). This experimental confirmation solidified the inhibitory capabilities of WMiApt-1 in blocking WDR5-MYC interactions in a cellular context. Previous studies have demonstrated that disrupting the interaction between WDR5 and MYC can arrest MYC-driven tumorigenesis^57^. Indeed, assessing its impact on cell proliferation in the MDA-MB-468 cell line revealed the effectiveness of WMiApt-1 in arresting cell proliferation (Supplementary Fig. 9s), but further investigations, which are outside the scope of this paper, are required to elucidate the underlying mechanism. These findings affirm the potential of WMiApt-1 as a valuable chemical tool for targeting WDR5-MYC interaction.

Collectively, our proposed structure-based design strategy, Blocker-SELEX, has facilitated the successful development of inhibitory aptamers capable of disrupting interactions involving transcription factors, thereby opening new avenues for the exploration of inhibitory aptamer-based therapeutics.

## Discussion

Aptamers, distinguished by their several distinctive merits, have emerged as powerful chemical tools with broad applicability. However, the development of aptamers tailored to specific application scenarios remains a formidable challenge. Traditional aptamer-selection strategies, such as SELEX and Cell-SELEX^58^, are recognized for their labor-intensive and time-consuming nature, requiring multiple cycles of affinity selection to obtain high-affinity aptamers. Recent advancements have been made to enhance SELEX methodologies, including Pro-SELEX, which leverages microfluidic technology to enable the quantitative isolation of aptamers with programmable binding affinities^59^. Additionally, a functional group-guided approach has been proposed to enhance the selectivity of aptamers targeting small molecules^60^. Nonetheless, a significant research gap exists in the development of selection methods explicitly designed to enable aptamers to bind to discrete protein regions of interest and modulate protein-protein interactions.

In this study, we introduce Blocker-SELEX, a structure-guided aptamer design pipeline. This approach efficiently generates competitive iAptamer<u>s</u> that selectively bind to desired target interfaces, effectively blocking the associated interactions, and they do so without relying on computationally intensive processes or accurate 3D structure predictions. Blocker-SELEX assesses the binding potential of individual sequences to the target interface, enabling the scoring of competitive capabilities for each aptamer candidate in terms of target interactions. Moreover, Blocker-SELEX serves as a valuable starting point for constructing potent aptamers within specific binding regions. This reverse process of traditional aptamer SELEX effectively addresses the challenge of identifying essential and nonessential regions of aptamers post-SELEX, thereby facilitating the development of aptamers with enhanced functionality and specificity^61,62^.

By employing this strategy, we have effectively engineered inhibitory aptamers that possess the ability to disrupt the interactions between RNAP2 and SCAF4 or SCAF8 proteins, which were previously considered “undruggable” using small-molecule inhibitors owing to the lack of “conventional drug-binding” pockets at the interface of SCAF4/SCAF8. Notably, aptamers exhibit target recognition capabilities that do not necessitate the presence of a “conventional drug-binding” pocket^43,63–68^. This characteristic makes aptamers ideal chemical tools for intervening in “undruggable” protein-protein/DNA interactions. By utilizing this approach, we validated the feasibility of structure-guided aptamer design and screening. Given the advancements in artificial intelligence within the structural domain and the increasing computational power, it is now foreseeable that we will be capable of *de novo* designing longer nucleic acid aptamers based on their structural attributes.

Overall, Blocker-SELEX offers a significant advantage in designing functional aptamers that can target specific regions of a protein with distinct functions, encompassing the blocking of protein-protein interactions, transcription factor-DNA/RNA interactions, and even extending to the inhibition of enzyme-substrate interactions or acting as allosteric modulators. This unique capability enables the precise regulation of functional signal pathways through the utilization of aptamer inhibitors, thereby presenting a promising avenue for advancements in drug discovery. By harnessing the full potential of the Blocker-SELEX pipeline, our objective is to revolutionize aptamer design and facilitate the creation of tailor-made aptamers for precise applications, thereby expanding their versatility across diverse fields.

## Methods

### Virtual screening

A library containing 1024 ssDNA sequences with a length of 5 nucleotides was prepared by JChem software (http://www.chemaxon.com) and optimized using the LigPrep module from Schrodinger suite software (release 2019-02)^46^. Atomic protonation was adjusted to pH 7.0 with the Epik software^69^, and geometric optimization was performed using the OPLS_2005 Force Field software^70^. Protein structures used for docking were retrieved from the Protein Data Bank (SCAF4: 6XKB; SCAF8: 3D90 chain B; WDR5: 4Y7R chain A) (https://www.rcsb.org/). Protein preparation was performed with the Protein Preparation Wizard from the Schrödinger platform using default settings to remove all crystallographic water molecules, add hydrogen atoms, assign partial charges, and minimize structures^46^. Targeted docking was carried out on SCAF4, SCAF8 or WDR5 proteins. Grid boxes were set to 25 Å × 25 Å × 25 Å for ligand docking centered at each bound ligand by using the receptor grid generation pipeline^48^. Molecular docking was performed with the ligand docking pipeline with ligand sampling set to flexible in a standard precision scale^71^.

### Protein expression and purification

The Human SCAF4-CID domain fragment comprising residues 1-139 and SCAF8-CID domain fragment comprising residues 1-139 were subcloned into a modified pET28-MHL vector, generating N-terminal His-tagged fusion proteins. The hSCAF4-CID mutants were obtained via Quick site-directed mutagenesis utilizing the hSCAF4-CID (residues 1-139) expression construct as the template. The recombinant protein was expressed in BL21 (DE3) cells and induced with 0.5 mM isopropyl-β-D-thiogalactopyranoside (IPTG) at 16°C overnight. Cells were collected via centrifugation, resuspended in lysis buffer containing DPBS buffer, 0.05 mM EDTA, and 5 mM imidazole, and sonicated. The supernatant was obtained after centrifugation at 9500g for 1 hour and further purified via the Ni-NTA Beads Gravity Column (Changzhou Smart-Lifesciences Biotechnology Co., China). His-tag was cleaved using TEV Protease, and the protein was subsequently purified using a Superdex75 gel-filtration column (GE Healthcare, Chicago, IL, USA). Finally, purified proteins were concentrated to 2 mg/mL in DPBS buffer supplemented with 5% glycerol. WDR5 proteins were purified, as previously described^72^. Briefly, 6xHis-SUMO-tagged WDR5 protein was expressed in BL21 (DE3) cells upon reaching an optical density of OD = 0.8 via induction with 0.5 mM IPTG at 16 ℃ for 18 hours. Recombinant WDR5 protein was purified to homogeneity using Ni-NTA. The His-SUMO-tag was removed by Thrombin protease cleavage during dialysis and subsequent subtractive second nickel-column^73,74^.

### Surface Plasmon Resonance Experiments

Lead sequence screening: Identification of lead sequences was carried out using surface plasmon resonance (SPR) analysis, employing a BIAcore 8K instrument with CM5 chips (GE Healthcare) at ambient temperature (25 ℃). All proteins used in SPR analysis were exchanged to DPBS buffer (2.68 mM KCl, 1.47 mM KH_2_PO_4_, 136.89 mM NaCl, 8.09 mM Na_2_HPO_4_·12H_2_O, pH 7.5) containing 0.002% (v/v) Tween-20. Aptamer candidates were diluted to a concentration of 10 μM and then passed over the chip surface to measure response units. Aptamers exhibiting high response levels were deemed as lead sequences.

Affinity determination: Measurement of binding affinity was carried out using SPR analysis with a BIAcore 8K instrument and CM5 chips (GE Healthcare) at ambient temperature (25 ℃). Aptamers were diluted serially to a range of concentrations, and the analytes were passed over the chip surface to measure response units. Binding kinetics was analyzed using the Biacore Insight Evaluation Software with a 1:1 Langmuir binding model. For interactions between aptamers and SCAF4/8 proteins, the SPR buffer consisted of DPBS supplemented with 0.002% (v/v) Tween-20. In the case of interactions between aptamers and the WDR5 protein, the SPR buffer comprised MES buffer (30 mM MES buffer, pH 6.5, 25 mM NaCl, 2 mM β-ME, 1 mM CHAPS, and 0.002 mg/mL BSA).

### Fluorescence polarization

Experimental procedures were carried out using a Perkin Elmer EnVision® 2104 Multilabel Reader (BioTek, USA) equipped with 485□nm excitation and 535□nm emission filters for the FITC. FP measurements were performed using a Bioland 96-well plate (product #PB06-96S). Each well was loaded with 100□μL assay solution containing FITC-labeled peptides or aptamers (FITC-labeled, 50□μL) in the concentration of 50□nM and proteins (50□μL). Protein samples were serially diluted to various concentrations. After a 30-minute incubation period at room temperature, FP measurements were taken. Both parallel and perpendicular fluorescence intensity (F□ and F⊥) relative to linearly polarized excitation light were determined to calculate the FP signal. The experiments were conducted in triplicate, and the average affinity value was determined using the GraphPad Prism 9.0.0 program (GraphPad Software, Inc., USA) through curve fitting.

For the interactions of SCAF4-RNAP2 peptide or SCAF4-Aptamers, the FP assay buffer was constituted by DPBS buffer (2.68 mM KCl, 1.47 mM KH_2_PO_4_, 136.89 mM NaCl, 8.09 mM Na_2_HPO_4_·12H_2_O, pH 6.5), 2 mM β-ME, 1 mM CHAPS, and 0.002 mg/mL BSA.

For the interaction between WDR5 and MYC peptide, the FP assay buffer was constituted by 30 mM MES buffer pH 6.0, 25 mM NaCl, 2 mM β-ME, 1 mM CHAPS, and 0.002 mg/mL BSA.

For the interaction between WDR5 and Aptamers, the FP assay buffer was constituted by 30 mM MES buffer pH 6.5, 25 mM NaCl, 2 mM β-ME, 1 mM CHAPS, and 0.002 mg/mL BSA.

### Simulation of L2-SCAF4 complex

The docked complex of SCAF4 and SCAF4_LS was used as the initial starting point for the MD (Molecular Dynamics) simulation study for assessing their stability as a complex in terms of Root Mean Square Deviation (RMSD), Root Mean Square Fluctuation (RMSF) and other parameters. MD Simulation was performed by Gromacs 2021.7 software^75^. CHARMM36m force-field was used for protein^76^ and DNA^77^ in complex. The initial binding conformation was obtained via template-guided molecular docking using S2, S5-quadra-phosphorylated CTD peptide as the template (PDBID:6XKB). Solvent water, as charmm-tip3p water model, and ions (Na^+^, K^+^ and HPO_4_ ^2-^) were added via Gromacs to emulate PBS buffer. After energy minimization, temperature of the system was heated from 0 K to 298 K via a 500 ps annealing simulation (NPT ensemble) with position restraints on the backbone of protein and DNA. Then, final equilibrium NPT ensemble simulation of 1 ns without any restraint at 298 K was done before production phase simulation. A 500 ns NPT simulation was conducted after equilibrium, and the last 100 ns of trajectory were extracted for post-simulation analysis. RMDS of DNA in the trajectory was calculated by superimposing the protein and comparing the result to the final conformation. The RMSF was calculated in the same way, but compared to the average conformation. Temperature and pressure of the system were controlled by Bussi-Parinello (Stochastic) Velocity Rescaling^78^ and Stochastic Cell Rescaling^75^ during all MD simulation processes.

### Simulation of WMiApt-1-WDR5 complex

The HADDOCK server^79^ was used to generate the complex structure of WDR5 and WMiApt-1 by loading the cleaned WDR5 structure (PDBID: 4R7Y) and WMiApt-1 to the web server and docking with standard parameters through a hybrid algorithm of template-based and template-free docking. The docked complex structure were used as the initial starting point for the MD (Molecular Dynamics) simulation study for assessing their stability in terms of Root Mean Square Deviation (RMSD), Root Mean Square Fluctuation (RMSF) and other parameters. MD Simulation was performed using the same protocol as L2-SCAF4 complex simulation. Due to the large complex WDR5-WMiApt, we have performed 1 μs of simulation and the last 200 ns of trajectory was extracted.

### Competition assays of inhibitory aptamers

Competition assays of inhibitory aptamers for protein-protein interaction were performed using Fluorescence Polarization (FP) assays. In brief, FP signals resulting from the interaction were detected by combining 50□nM of FITC-labeled peptides with SCAF4/8 or WDR5 proteins in the FP competition buffer (30mM MES pH 6.0, 25mM NaCl, 2mM β-ME, 1mM CHAPS, and 0.002mg/mL BSA). Each well of a 96-well plate was loaded with 100□μL of the assay solution. Aptamers were serially diluted to varying concentrations and added to each well for the competition assay. Experiments were performed in triplicate, and the average inhibition constant values were determined by performing curve fitting and data analysis using GraphPad Prism 9.0.0 (GraphPad Software, Inc., USA).

### Inhibitory aptamer screening by competition assay

Experiments were conducted using a Perkin Elmer EnVision® 2104 Multilabel Reader (BioTek, USA) with 485□nm excitation and 535□nm emission filters for the FITC. The competition assay was performed using a Bioland 96-well plate (product #PB06-96S). To generate the FP signal, 50□nM FITC-labeled peptides or aptamers were mixed with 1 μM target proteins and plated into wells of a 96-well plate at a volume of 100 μL per well. The FP signal was measured for each well, and optimized aptamers were introduced into each well for the competition assay. The FP signal was measured again, and reduction of the FP signal was calculated. Aptamers that exhibited the most significant FP signal reduction were regarded as highly competitive.

Curve fitting and data analysis were performed using GraphPad Prism 9.0.0 (GraphPad Software, Inc., USA). Experiments were performed in triplicate, and the mean results were reported.

For SCAF4 assays, the assay buffer was composed of DPBS buffer supplemented with 2 mM β-ME, 1 mM CHAPS, and 0.002 mg/mL BSA.

For WDR5 assays, the assay buffer was constituted by 30 mM MES buffer pH 6.0, 25 mM NaCl, 2 mM β-ME, 1 mM CHAPS, and 0.002 mg/mL BSA.

### Immunoprecipitation

Immunoprecipitation (IP) was performed following the manufacturer’s instructions using the BeyoMag™ Streptavidin Magnetic Beads IP Kit. Biotinylated aptamers, or reverse complementary sequences (30 μg), were mixed with 20□μL of BeyoMag™ Streptavidin Magnetic Beads in a tube containing 20□μL of PBST and incubated at room temperature for 2 hours. Biotinylated SCAF4_F3_11nt aptamers or NC-loaded beads were then obtained by removing the supernatant on a magnetic separator. To prepare whole cell extracts, cell pellets were lysed using NP-40 lysis buffer containing 50 mM Tris-HCl pH 7.5, 500 mM NaCl, 2 mM EDTA, 0.5% (v/v) Triton X-100, PhosSTOP (Roche, 04906837001) and Protease Inhibitor Cocktail (Roche, 05056489001). Samples were centrifuged, and the supernatants were mixed with the biotinylated immunomagnetic beads to form an immunomagnetic beads-aptamer-antigen complex. Beads were washed three times with PBS and subjected to Western blotting analysis.

Cells transfected with indicated plasmids (SCAF4-CID-3xHA-pCDNA3.1 or MYC-pCDNA3.1) were lysed in cell lysis buffer (150□mM NaCl, 10□mM Tris/HCl pH 7.4, 1□mM EDTA, 1□mM EGTA, 0.2% NP-40, and protease inhibitor mixture). The cell lysate was treated with PBS buffer (control) or inhibitory aptamers in 10 µM and incubated with anti-HA antibody (Abclonal, AE008, Mouse) or anti-MYC antibody (Sangon, D199941-0100, Mouse) overnight at 4 ℃. Normally, 600□µg of total proteins were used for Co-IP in a total volume of 700□µL. Protein A/G conjugated beads were added to precipitate the antibodies. After removing the supernatant and washing with 700□µL of lysis buffer five times, the immunoprecipitates were processed for Western blotting.

### Western blotting

Proteins captured on beads were solubilized by adding SDS-loading buffer, subjecting to electrophoresis, and transferring onto a Polyvinylidene Fluoride (PVDF) membrane (Millipore, ISEQ00010). Membranes were subsequently incubated with the appropriate primary antibodies: anti-SCAF4 (Sangon, D153368-0100, Rabbit), anti-SCAF8 (Abclonal, A19467, Rabbit), anti-RNAP2 (Abclonal, A2107, Rabbit), anti-Phospho-RNAP2-S5 (Abclonal, AP0828, Rabbit), anti-WDR5 (Abclonal, A3259, Rabbit), and anti-MYC antibody (Abclonal, A1309, Rabbit), followed by incubation with horseradish peroxidase (HRP)-conjugated secondary antibodies (WanleiBio, China). Proteins of interest were visualized using an enhanced chemiluminescence detection kit (Biosharp, China).

### Aptamer-LNPs assembly

The preparation of Aptamer-LNPs was carried out using the ethanol dilution method described elsewhere^80^, using DOTAP, D-Lin-MC3-DMA, DOPE, cholesterol and DMG-PEG2k as the reagents with molar ratio of 50: 25: 5: 19.3: 0.8. Final mass ratio of lipid material to Aptamer was 40:1, and the volume ratio of ethanol to aqueous was 1:3. The assembled LNPs were dialyzed in 1× PBS using a 3500 Da dialysis bag for 2 hours, followed by centrifugation at 5000 rpm at 4 ℃. The concentrated Aptamer-LNPs were then stored in the dark at 4 ℃. In the same way, NC-LNPs were prepared by replacing AM with a negative control sequence. The preparation of SM-102-based-LNPs (Moderna’s LNP formulation) was carried out as described^81^.

### Cell proliferation assay

Human colon cancer cell line HCT116 was cultured in RPMI-1640 (YESEN, 41402ES76) with 10% FBS (Sunrise, SR100180.03) and 1% penicillin/streptomycin (Biosharp) and incubated in a 5% CO2 incubator at 37 ℃. For inhibitory aptamer treatment, cells were seeded into a 96-well plate at a density of 15K cells per well and exposed to 1 μM SRiApt aptamer or NC packed in LNPs for 5 days. Cell viability was determined by Cell Counting Kit-8 assay (Biosharp, BS350B).

### Apoptosis

HCT116 cells (15K cells/well) were seeded in wells of a 96-well plate and treated with 1 μM SCAF4_F3_11nt aptamer or NC packed in LNPs for 3 days. Annexin V reagent (Vazyme) was added and analyzed with the Incucyte SX5 live cell imaging device (Sartorius, Germany).

### Confocal imaging

HCT116 cells were seeded and treated with SRiApt-1 or SRiApt-3 aptamer-SM102 in a final concentration of 1 μM for 6 hours. After treatment, cells were washed with PBS and then fixed in 4% multigrade formaldehyde (Biosharp) for 1 hour at room temperature. Cells were then washed with PBS and blocked in 5% bovine serum albumin (Biosharp) for 1 hour at room temperature. Cells were washed with PBS, and DAPI (Beyotime) was added and incubated for 1 minute. Images were detected on a single-photon confocal microscope (NIKON, A1 HD25) and analyzed by ImageJ (v1.8.0).

### RNA-seq analysis

HCT116 cells were subjected to treatment with SRiApt-1 aptamers or NC at a final concentration of 1 µM for 36 hours, followed by collection for Bulk RNA-seq. Library construction and sequencing were conducted by GENEWIZ (GENEWIZ Biotechnology Co., Ltd., Suzhou, China) on an Illumina HiSeq instrument with a 2×150 paired-end (PE) configuration in accordance with the manufacturer’s instructions. The reference genome sequences were indexed using Hisat2 (v2.0.1), and clean data were aligned to the reference genome using the same software. The known gff annotation files were converted into transcripts in fasta format and indexed appropriately. With the reference gene file, HTSeq (v0.6.1) estimated gene and isoform expression levels from the pair-end clean data. The DESeq2 package (https://github.com/mikelove/DESeq2) was used to perform gene expression analysis and rank the results based on stat value. The Hallmark and KEGG gene sets were obtained from MsigDB (https://www.gsea-msigdb.org/gsea/msigdb/). The clusterProfiler R package was used to perform GSEA (gene set enrichment analysis), and the enrichplot R package was employed to generate pathway enrichment plots.

### Alternative splicing analysis

The input for the analysis consisted of RNA-seq generated bam files aligned using the HISAT2 aligner. The computational tool rMATs v4.1.0 was employed to identify differential alternative splicing events from the RNA-seq data. To ensure high confidence, all alternative splicing events were filtered based on an FDR of ≤0.05 and an inclusion level difference of > 0.02. The number of events per comparison was then calculated.

## Supplementary Materials

Supplementary Materials are available online.

## Acknowledgements

We thank the model of organized joint research of HIM. This work was supported by grants from the National Key R&D Program of China (2022YFC3400400 to Y.W.), National Natural Science Foundation of China (No. 32201010, No. 82103287, T2188102), Zhejiang Provincial Natural Science Foundation of China (grant LR22B050001 to Q.W., YXD24B0701 to Y.W.), Zhejiang Provincial Research Center for Diagnosis and Treatment of Major Diseases (JBZX-202003), Zhejiang Provincial Natural Science Foundation (grants LR22B050001, Y21C050001), and Zhejiang Province Postdoctoral Research Project Merit-based Funding Project (ZJ2022048 to X. Liu).

## Author contributions

W. Tan, Q. Wu conceived the project. T. Li, X. Liu performed the bulk of experiments. Y. Wei, Y. Tao performed virtual screening. Y. Zhang, C. Sha performed Molecular Simulation and Modeling. A. Lin, T. Li performed the assembly of Aptamer-LNPs. J. Qin, W. Chen, Y. Hou, G. Luo, X. Zhu helped. X. Liu performed the bioinformatics analysis. Y. Wei, X. Liu wrote the manuscript. W. Tan, Q. Wu supervised the work. All authors discussed the results and contributed to the final manuscript.

## Conflict of Interests

The authors declare no competing interests.

## Notes

### Competing Interest Statement

The authors have declared no competing interest.

